# Engineered developmental niche enables predictive phenotypic screening in human dystrophic cardiomyopathy

**DOI:** 10.1101/456301

**Authors:** Jesse R. Macadangdang, Jason W. Miklas, Alec S.T. Smith, Eunpyo Choi, Winnie Leung, Yuliang Wang, Xuan Guan, Soowan Lee, Max R. Salick, Michael Regnier, David Mack, Martin K. Childers, Hannele Ruohola-Baker, Deok-Ho Kim

**Affiliations:** Bioengineering, University of Washington, Seattle, WA; Center for Cardiovascular Biology, University of Washington, Seattle, WA; Institute for Stem Cell and Regenerative Medicine, University of Washington, Seattle, WA; Rehabilitation Medicine, University of Washington, Seattle, WA; Biochemistry, University of Washington, Seattle, WA; Computer Science and Engineering; Department of Engineering Physics, University of Wisconsin, Madison, WI; Wisconsin Institute of Discovery, Madison, WI

**Author notes:** These authors contributed equally (Jesse R. Macadangdang, Jason W. Miklas). Corresponding Authors Deok-Ho Kim, Ph.D., University of Washington, Department of Bioengineering, N410G William H Foege Building, 3720 15^th^ Ave NE, Box 355061, Seattle, WA 98195, Hannele Ruohola-Baker, Ph.D., University of Washington, Department of Biochemistry, S580 UW Medicine at South Lake Union, 850 Republican Street, Box 358056, Seattle, WA 98109.

## Abstract

Directed differentiation of human pluripotent stem cells (hPSCs) into cardiomyocytes typically produces cells with structural, functional, and biochemical properties that most closely resemble those present in the fetal heart. Here we establish an *in vitro* engineered developmental cardiac niche to produce matured hPSC-derived cardiomyocytes (hPSC-CMs) with enhanced sarcomere development, electrophysiology, contractile function, mitochondrial capacity, and a more mature transcriptome. When this developmental cardiac niche was applied to dystrophin mutant hPSC-CMs, a robust disease phenotype emerged, which was not observed in non-matured diseased hPSC-CMs. Matured dystrophin mutant hPSC-CMs exhibited a greater propensity for arrhythmia as measured via beat rate variability, most likely due to higher resting cytosolic calcium content. Using a custom nanopatterned microelectrode array platform to screen functional output in hPSC-CMs exposed to our engineered developmental cardiac niche, we identified calcium channel blocker, nitrendipine, mitigated hPSC-CM arrhythmogenic behavior and correctly identified sildenafil as a false positive. Taken together, we demonstrate our developmental cardiac niche platform enables robust hPSC-CM maturation allowing for more accurate disease modeling and predictive drug screening.

## INTRODUCTION

Cardiovascular disease remains the leading cause of death for both men and women worldwide, with a rapidly growing impact on developing nations^1^. Inherited cardiomyopathies are the major cause of heart disease in all age groups, including children and young, otherwise healthy adults^2^. Human pluripotent stem cell-derived cardiomyocytes (hPSC-CMs) are a promising tool for studying cardiomyopathy because they vastly increase the accessibility to human cardiomyocytes that can theoretically express the full array of ion channels, protein isoforms, and genetic and metabolic machinery found in the human heart^3^. A major goal of current cardiac disease modeling efforts is to gain insight into the onset and progression of cardiomyopathies as a first step towards designing more effective therapeutic strategies. Many patients with inherited cardiomyopathies, however, do not present with cardiac symptoms until late adolescence or early adulthood^2^. This limits the potential of hPSC-CM disease modeling since current differentiation protocols produce cardiomyocytes with structural, functional, and biochemical properties that most closely resemble cells of the fetal heart^4^.

Developmental maturation presents a technical hurdle for hPSC-CMs in cardiac disease models. To date, hPSC-CMs exhibiting a fetal-like phenotype have been used to study channelopathies^5^^-^^7^, metabolic syndromes^8^, hypertrophic and dilated cardiomyopathies^9^^,^^10^, as well as other delayed-onset cardiac diseases such as Duchenne Muscular Dystrophy (DMD) cardiomyopathy^11^^,^^12^. While such reports have certainly been groundbreaking and valuable^13^, the majority of the findings are based on the performance of cardiomyocytes that are still quite naive, having incomplete structural organization, and a mixture of fetal and adult protein isoform expression. Much effort has been invested to develop methods to accelerate the development of hPSC-CMs^4^^,^^13^. While such strategies have proven to be effective at enhancing specific aspects of hPSC-CM development, it is unlikely that a single method can fully recreate the developing cardiac microenvironmental niche. We therefore generated a multifaceted and combinatorial maturation (ComboMat) strategy, incorporating nanotopographic substrate cues, thyroid hormone T3, and Let-7i miRNA overexpression aimed at more comprehensively mimicking the developmental cardiac niche in order to accelerate the maturation of hPSC-CMs *in vitro*. Anisotropic extracellular matrix cues, critical for proper structure-function relationship in the fully developed heart and morphological development in hPSC-CMs^14^, have also been found to be an important factor during cardiac development and looping^15^. Thyroid hormone treatment, simulating the spike in serum T3 levels immediately following birth, has been shown to improve force production and calcium handling in hPSC-CMs^16^, while upregulation of Let-7 miRNAs enhance metabolic development in hPSC-CMs^17^.

As a means to highlight the utility of a mature cardiac phenotype for *in vitro* cardiac tissue engineering applications, we set out to develop a more mature model of dystrophic cardiomyopathy. Dystrophic cardiomyopathies, such as Duchenne muscular dystrophy (DMD) and Becker muscular dystrophy (BMD), are X-linked genetic disorders resulting from a mutated dystrophin gene. Cardiac complications have replaced respiratory failure as the leading cause of death in DMD patients due to the emergence of supportive ventilation as a standard of care^18^. However, cardiomyopathy in dystrophic rodent models, such as the *mdx* mouse, have proven to be poor predictors of patients' responses to therapeutic treatment in the clinic^19^^,^^20^. In fact, a recent clinical trial with sildenafil, which showed promise in alleviating cardiac dysfunction in the *mdx* mouse^21^, exacerbated cardiac symptoms in DMD patients and had to be terminated early^20^. Thus, an unmet medical need exists for improved human models of dystrophic cardiomyopathy. We previously reported that dystrophin mutant human cardiomyocytes displayed phenotypic differences compared to healthy controls, but these differences were mild or were induced by exogenous acute stress, such as hypotonic challenge^11,22^. Other dystrophin mutant models, developed by Lin et al.^12^ and Young et al.^23^, provide valuable insight into DMD-specific disease mechanisms and potential therapeutic treatment options, but are based on fetal-like hPSC-CMs. We hypothesize that more mature hPSC-CMs will improve the predictive power of preclinical screening models of dystrophic cardiomyopathy and create an *in vitro* phenotype more similar to that found in patients.

To test this hypothesis, we applied our ComboMat strategy to CRISPR-generated dystrophin mutant (*DMD* 263delG) hPSC-CMs in order to facilitate the manifestation of adult-onset cardiac disease phenotypes *in vitro*. Patients with similar exon 1 mutations in the *DMD* gene (**Supplementary Fig. 1**) express a truncated dystrophin protein and present with a BMD phenotype^24^. When matured using our combinatorial maturation strategy, *DMD* mutant hPSC-CMs exhibit a much greater propensity for arrhythmia, as measured using our custom nanopatterned multielectrode array (nanoMEA) cardiac screening platform. Without developmental cardiac niche cues, we could not distinguish the functional profile of *DMD* mutant hPSC-CMs from healthy isogenic control cells. Thus, the ComboMat platform produced more physiologically relevant hPSC-CMs for disease modeling and drug screening applications.

## RESULTS

### Profiling the transcriptome of ComboMat-treated cardiomyocytes

We investigated the combinatorial impact of three distinct cues, biomimetic nanopatterned topography (NP)^14^^,^^15^, thyroid hormone T3 (T3)^16^, and Let7i miRNA overexpression (Let7i)^17^, incorporated into a single procedure termed the ComboMat platform (schematically depicted in **Fig. 1a**). We performed RNA-seq to look at the whole transcriptome of hPSC-CMs exposed to different combinations of the three maturation cues. Principal component analysis (**Fig. 1b**) demonstrated that the ComboMat group (all three factors combined together) separated from all other conditions. Next, we generated an enrichment heat map that consists of seven hallmark pathways of cardiomyocyte maturation, (**Fig. 1c**). As more maturation conditions were layered on top of one another, cardiomyocytes appeared to be progressively more mature. The ComboMat group showed the most robust maturation, with up-regulation in six of the pathways and down-regulation in the cell cycle pathway. **Figure 1d** shows 8 up-regulated gene ontology (GO) terms based on *P*-value between the ComboMat and Control (Empty Vector cells on flat surfaces; EV-Flat) groups. Pathways related to metabolism such as glucose metabolic processes, long-chain fatty acid import, and glycolytic process were up-regulated. GO terms associated with muscle processes including muscle tissue development, muscle contraction, and striated muscle cell differentiation were also up-regulated.

**Figure 1.**
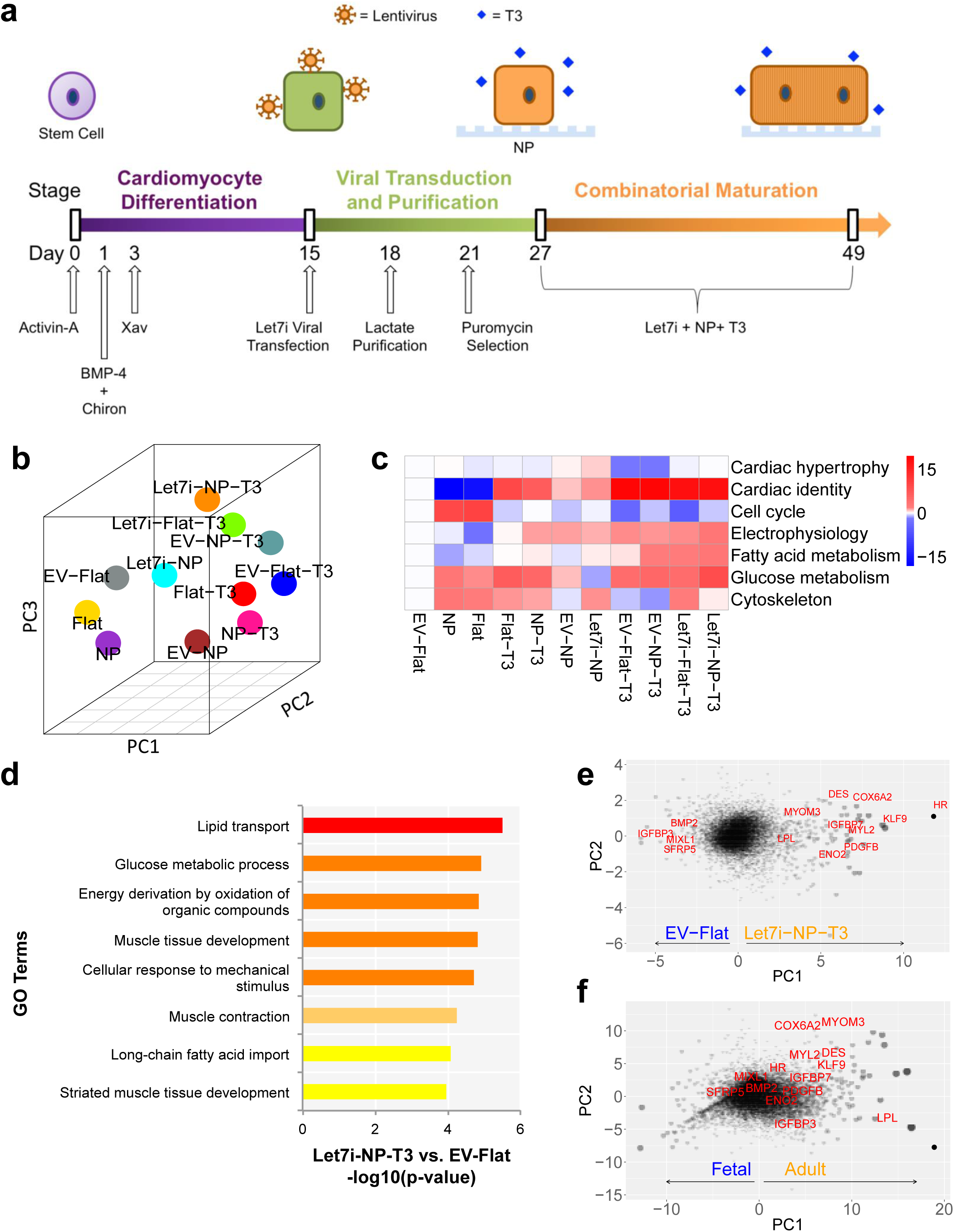
ComboMat platform promotes a more mature transcriptional profile. (**a**) Schematic timeline of the ComboMat platform. Cardiomyocyte differentiation is achieved using a small molecule and human recombinant protein-based protocol. After cardiomyocyte production, hPSC-CMs are transfected with Let7i lentivirus, purified via metabolic challenge, and selected for viral transfection. High purity hPSC-CMs that express the viral vector are then plated on NPs and exposed to T3 for 3 weeks. (**b**) Visualization of the Principle Component Analysis in three dimensions. hPSC-CMs exposed to T3 cluster along PC2. The ComboMat group separates and is distinct from other combinations of the maturation stimuli. (**c**) Net enrichment in cardiac maturation pathways of differentially expressed genes as a result of different treatments. Red indicates genes up-regulated by the treatment show more significant enrichment in the pathway than genes downregulated by the treatment; blue means the opposite. (**d**) Gene ontology enrichment results of up-regulated genes in Let7i-NP-T3 combination treatment compared to EV-Flat control. Color gradient and bar lengths represent significance of enrichment. (**e,f**) Bubble plot of PCA loadings showing genes’ contributions to the separation of RNAseq samples comparing EV-Flat to Let7i-NP-T3 (**e**) and fetal to adult cardiac cardiomyocytes (**f**). Bubble plots in (**e**) and (**f**) highlight a similar up/down-regulation pattern of specific genes between the ComboMat-treated hPSC-CMs and adult CMs.

We generated a bubble plot (**Fig. 1e**) to compare gene expression of the ComboMat and Control groups along PC1. We found 53 genes that were significantly higher and 13 genes that were significantly lower in both Adult vs. Fetal and ComboMat vs. Control. The gene that was most up-regulated along PC1 in the ComboMat group was Hair Growth Associated gene (HR), a thyroid hormone co-repressor. Krupel-like factor (KLF9), a transcription factor, was also significantly up-regulated. KLF9 has been reported to bind to the PPARγ promoter, a critical gene in fatty acid metabolism^25^. Cytochrome c oxidase subunit 6A2 (COX6A2), another gene associated with cardiac metabolism, was also up-regulated in the ComboMat group. Furthermore, myosin light chain 2 (MYL2), the ventricular isoform and a hallmark of cardiomyocyte maturation, was up-regulated. A bubble plot displays human fetal and adult cardiomyocyte gene expression (**Fig. 1f**) compared to the same genes highlighted in **Figure 1e**. The highly up-regulated genes in the ComboMat group are also up-regulated in adult cardiomyocytes when compared to fetal controls.

### Structural maturation of ComboMat-treated cardiomyocytes

For these studies, and all subsequent functional assays, we compared the full <u>ComboMat</u> platform (Let7i+NP+T3) to a viral vector <u>Control</u> (EV+Flat+No T3) and each maturation cue in isolation: <u>NP</u> (EV+NP+No T3), <u>T3</u> (EV+Flat+T3), and <u>Let7i</u> (Let7i+Flat+No T3). Immunofluorescence imaging (**Fig. 2a**) confirmed that hPSC-CMs cultured on NPs elongated in the direction of the nanotopography and exhibited more anisotropic, rod-shaped morphologies. Sarcomeres developed with a higher degree of directionality and order on NPs, resulting in more sarcomeres in register with one another (**Fig. 2b**). Compared to Control, hPSC-CMs exposed to just T3 or Let7i were significantly larger (**Fig. 2f**) but formed rounded or irregular morphologies. When exposed to all three cues in the ComboMat platform, hPSC-CMs became rod-shaped and were larger than with each cue in isolation (**Fig. 2f**, *p < 0.05). hPSC-CMs exposed to the ComboMat platform also displayed a repetitive sarcomere banding pattern along the length of the cell in contrast to the circumferential banding found in the Control group, and had longer resting sarcomere lengths of approximately 1.8 µm ± 0.012 µm (**Fig. 2c**, *p < 0.05). hPSC-CMs in the ComboMat group also exhibited a higher binucleation percentage (**Fig. 2e**, *p < 0.05).

**Figure 2.**
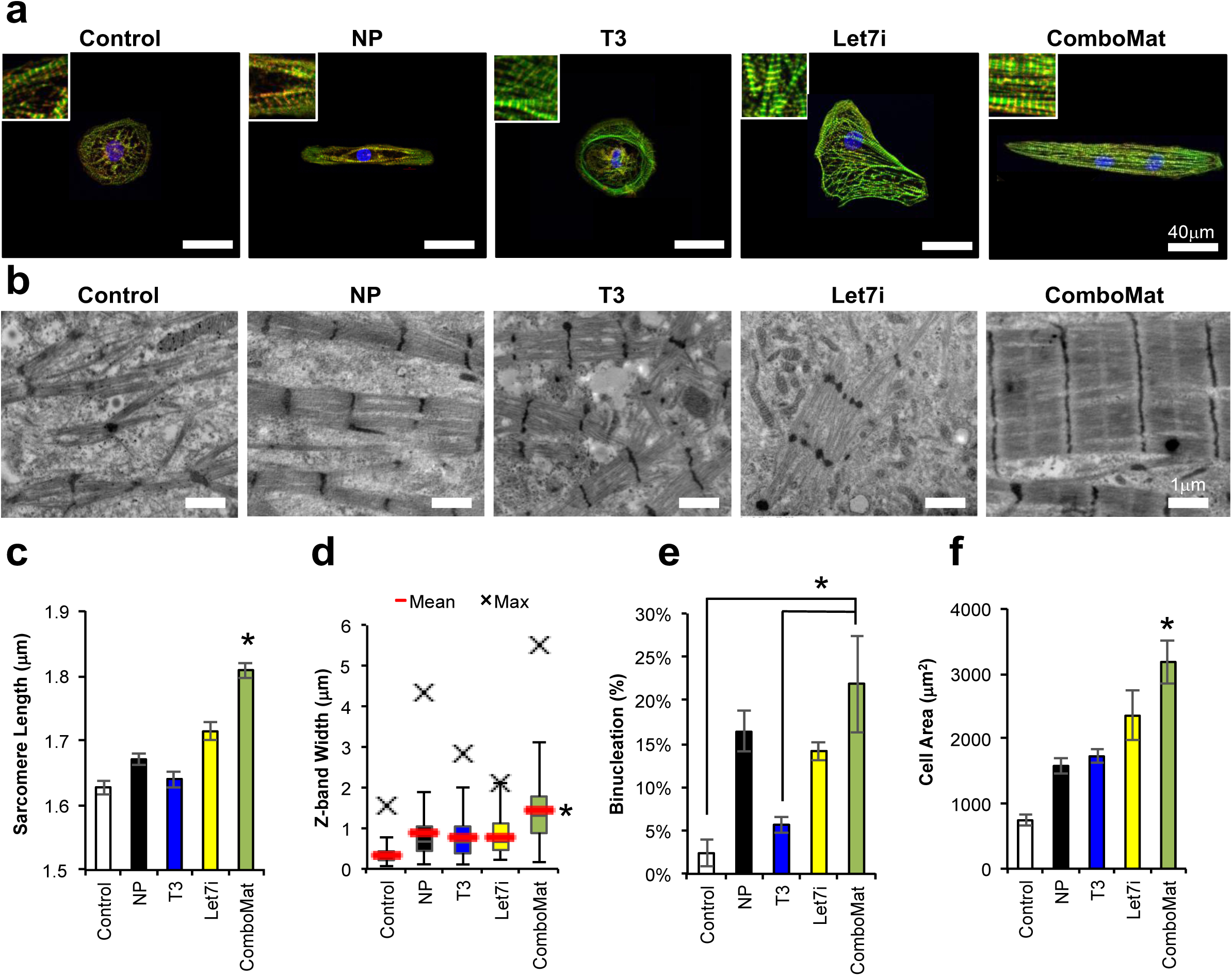
Structural maturation of hPSC-CMs. (**a**) Immunofluorescent images of hPSC-CMs exposed to the ComboMat platform or each cue in isolation. Red = α-actinin, Green = F-actin, Blue = DAPI. Insets show close-up of sarcomere structures within the hPSC-CMs. (**b**) TEM images of sarcomeres from hPSC-CMs exposed to the labeled cues of the ComboMat platform. hPSC-CMs exposed to the Control conditions exhibit only Z-body formation whereas hPSC-CMs exposed to the ComboMat platform have much more defined Z-disc structures. (**c**) Sarcomere lengths as measured via custom 2D Fourier transform image analysis. (**d**) Box-and-whisker plots of Z-band widths as measured from the TEM images with the mean Z-band width marked with the red line. (**e**) Bar chart of binucleated cells shows that the ComboMat group is approaching physiological binucleation percentages. (**f**) Cell area as measured from immunofluorescent images shows a clear hypertrophic response to the ComboMat platform. Bars represent averages ± S.E.M. *P<0.05.

The ultrastructure of the hPSC-CMs was investigated via transmission electron microscopy (TEM). We found that hPSC-CMs exposed to Control conditions exhibited low density, disorganized myofibrils and only punctate Z-body formation (**Fig. 2b**). The application of NPs, T3, or Let7i individually improved the development of more organized and wider Z-bands (**Fig. 2d**) but overall sarcomeres remained rather disorganized and at a low density. In contrast, hPSC-CMs cultured with the ComboMat platform developed much more ordered sarcomeres, with the emergence of Z-bands and H-zones (**Fig. 2b**). Z-band width in cells, an indicator of myofibril bundling, was also significantly larger in the ComboMat group than Control or each cue in isolation.

### Electro-mechanical and metabolic maturation of ComboMat-treated cardiomyocytes

To test if the genetic and structural changes imparted by the ComboMat platform manifested into enhanced functional performance of the hPSC-CMs, we employed two noninvasive measurements of electromechanical function. Using correlation-based contraction quantification (CCQ)^22^, we measured contractions of hPSC-CM monolayers paced at 1 Hz and filmed at 30 frames per second. By spatially averaging these contraction vectors together, we measured maximum contraction and relaxation velocities (annotated as red and blue marks in **Fig. 3a**) for the Control, NP, T3, Let7i, and ComboMat conditions and found that cells exposed to the ComboMat platform had significantly faster contractions compared to Control or any maturation cue in isolation (**Fig. 3b**). To help explain this, we examined the angles of displacement during contractions and found that hPSC-CMs cultured on NPs contract in a more uniform direction than cells cultured on flat substrates (**Fig. 3c**). To quantify this preferred directionality, we measured the contraction anisotropic ratio (AR) and found that cells on NPs, such as those in the ComboMat platform, had a contraction magnitude over two times greater in the direction parallel with the underlying nanotopography than perpendicular to it (ComboMat AR = 2.39 ± 0.01) (**Fig. 3d**). This was in contrast to cells grown on traditional flat substrates, where contractions were randomly oriented, and exhibited similar magnitudes in all directions (Control AR = 1.04 ± 0.02).

**Figure 3.**
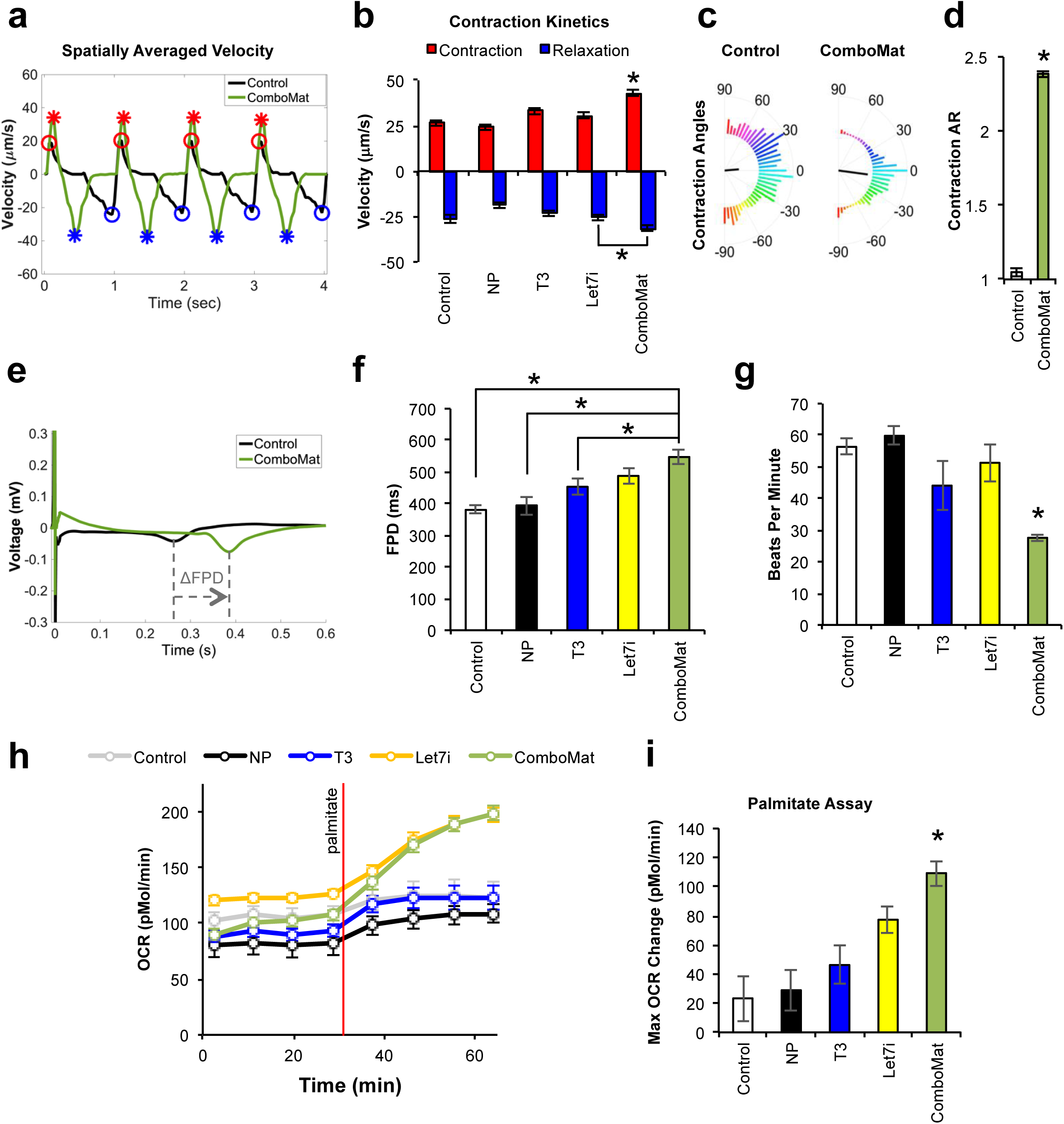
Electromechanical and metabolic maturation. (**a**) Representative traces of contraction velocities averaged across an entire video frame for Control or ComboMat hPSC-CMs as measured via our custom CCQ method. Red marks indicate peak contraction velocity, while blue demark peak relaxation velocity. (**b**) Quantification of peak contraction and relaxation velocities. (**c**) Contraction angle distribution histogram for Control or ComboMat hPSC-CMs. The black line in the center of the histogram denotes the average angle of contraction while the length of the black line indicates the degree to which the distribution is aligned with the average. (**d**) Contraction anisotropic ratio of the magnitude of contraction in the longitudinal and transverse directions. (**e**) Representative averaged field potential recording denoting the prolongation of the FPD between Control and ComboMat hPSC-CMs. Quantification of FPD (**f**) and beat rate expressed as Beats Per Minute (**g**) for hPSC-CMs exposed to the ComboMat platform or each cue in isolation. (**h**) Representative OCR trace for fatty acid stress test using palmitate. (**i**) Quantification of maximum change in OCR following palmitate addition. Bars represent averages ± S.E.M. *P<0.05.

In addition to contraction dynamics, we also measured cardiomyocyte electrical activity using microelectrode arrays (MEAs). An ion permeable resin, Nafion, was used to generate nanotopographic surfaces on MEAs (nanoMEAs) in order to facilitate hPSC-CM alignment in NP and ComboMat conditions while still enabling high resolution electrophysiological data capture from underlying electrodes. **Figure 3e** shows temporally averaged field potential recordings from Control and ComboMat cultures. Measuring spontaneous electrical activity of cardiac monolayers, we found that hPSC-CMs exposed to the ComboMat platform had significantly longer field potential durations than the Control, NP, and T3 groups (**Fig. 3f**). Additionally, the ComboMat platform significantly slowed the beat rate of the cardiomyocytes compared to Control or each maturation cue in isolation (**Fig. 3g**). hPSC-CMs exposed to the ComboMat platform also exhibited faster upstroke velocity as measured via patch clamp (**Supplementary Fig. 2**)

RNA-seq analysis suggested significant changes to cardiac metabolism imparted by the ComboMat platform. Thus, we probed the ability of the cardiomyocytes to utilize exogenous fatty acids via the “Palmitate assay” on a Seahorse mitochondrial flux analyzer (**Fig. 3h**). We found that hPSC-CMs exposed to the ComboMat platform exhibited significantly greater maximum change in oxygen consumption rate (OCR) measured between palmitate concentrations of 200 µM and 400 µM. (**Fig. 3i**). Maximum respiratory capacity was also significantly greater in the ComboMat group compared to Control (**Supplementary Fig. 3**)

### ComboMat-treated dystrophin-mutant hPSC-CMs manifest disease phenotype

To begin, we tested whether *DMD* mutant hPSC-CMs responded in a similar manner to the ComboMat platform as their Normal (UC3-4) isogenic counterparts. Using the Seahorse MitoStress assay (**Fig. 4a**), we found that *DMD* mutant hiPSC-CMs had an increased maximum respiratory capacity when cultured under the ComboMat platform compared to Control conditions (**Fig. 4b**). Within the Control or ComboMat conditions, however, there was no statistical difference between the Normal and *DMD* mutant groups. The baseline electrophysiological characteristics of the *DMD* mutant cells were measured using our custom nanoMEA platform. The beat rate of the *DMD* mutant hiPSC-CMs decreased in response to the ComboMat platform to the same extent as the Normal hPSC-CMs (**Fig. 4c**). Additionally, the field potential duration (FPD) of *DMD* mutant hPSC-CMs increased when exposed to ComboMat compared to Control, mirroring observations made from Normal hPSC-CMs (**Fig. 4d**). From a structural standpoint, the *DMD* mutant hiPSC-CMs developed a characteristic elongated morphology with in-register sarcomeres when exposed to the ComboMat platform, whereas *DMD* mutant hiPSC-CMs exposed to Control conditions were more rounded, with more disorganized sarcomeres (**Fig. 4e**).

**Figure 4.**
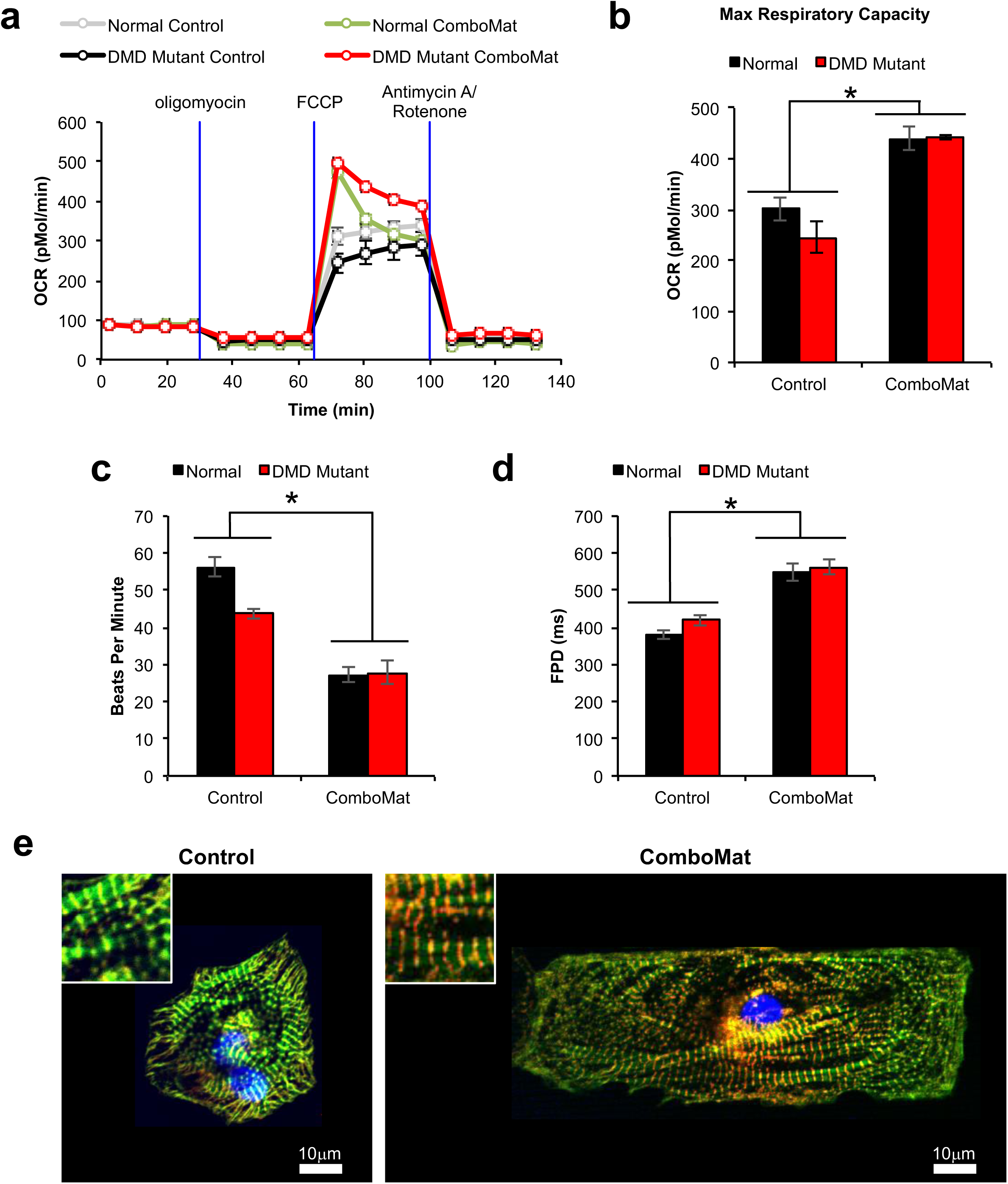
*DMD* mutant hPSC-CMs respond to ComboMat platform similarly as Normal control. (**a**) Seahorse metabolic profile of *DMD* mutant hPSC-CMs exposed to Control or ComboMat conditions compared to Normal hPSC-CMs. (**b**) Quantification of the maximum change in OCR after FCCP addition. Both Normal (black) and DMD mutant (red) hPSC-CMs show an increase in maximum respiratory capacity when exposed to the ComboMat platform. (**c,d**) MEA-based measurements of electrophysiology. (**c**) Beat rate represented as beats per minute for Normal or *DMD* mutant hPSC-CMs. (**d**) Average FPD for Normal or *DMD* mutant hPSC-CMs exposed to Control or ComboMat conditions. (**e**) Immunofluorescent images of *DMD* mutant hPSC-CMs exposed to Control (left) and ComboMat (right) conditions. Red = α-actinin, Green = F-actin, Blue = DAPI. Insets show zoomed-in portion of sarcomere structures in each condition. Bars represent averages ± S.E.M. *P<0.05.

We hypothesized that by inducing hypertrophy and overall cardiomyocyte maturation, dystrophic hPSC-CMs would undergo greater mechanical stress and cellular damage, resulting in irregular functional activity, which would be absent in immature cells. We used the nanoMEA platform to record the electrical activity of spontaneously beating monolayers of the isogenic pair of Normal and *DMD* mutant hPSC-CMs exposed to Control or ComboMat conditions. Using a method of arrhythmia detection relying on beat rate variability^26^, we found that *DMD* mutant hPSC-CMs exposed to the ComboMat platform were significantly more arrhythmogenic than less mature *DMD* mutant hPSC-CMs in the Control condition. **Figure 5a** shows representative field potential traces from *DMD* mutant hPSC-CMs exposed to Control (black) or ComboMat (red) conditions with individual beat intervals annotated. By measuring the difference in time of the beat interval from one beat to the next (ΔBI) and plotting this change for 30 consecutive beats (**Fig. 5b**), we found that matured *DMD* mutant hPSC-CMs had a striking instability in beat rate. By measuring the percentage of beats with a ΔBI > 250 ms (**Fig. 5c**) and the arithmetic mean for ΔBI over 30 consecutive beats (**Fig. 5d**), we saw relatively large changes in beat intervals indicative of a more arrhythmogenic state. Without an autonomous nervous system *in vitro*, hPSC-CMs should develop a steady beating pattern without much variation from beat to beat. This steady behavior is apparent in the Control conditions for both normal and *DMD* cardiomyocytes as well as in healthy mature Normal cells (**Fig. 5b**). Importantly, we found that the full ComboMat platform is necessary to observe functional differences between Normal and *DMD* mutant hPSC-CMs. When the maturation cues are applied in isolation, there is no separation between the healthy Normal cells and the *DMD* mutant hPSC-CMs (**Supplementary Fig. 4**)

**Figure 5.**
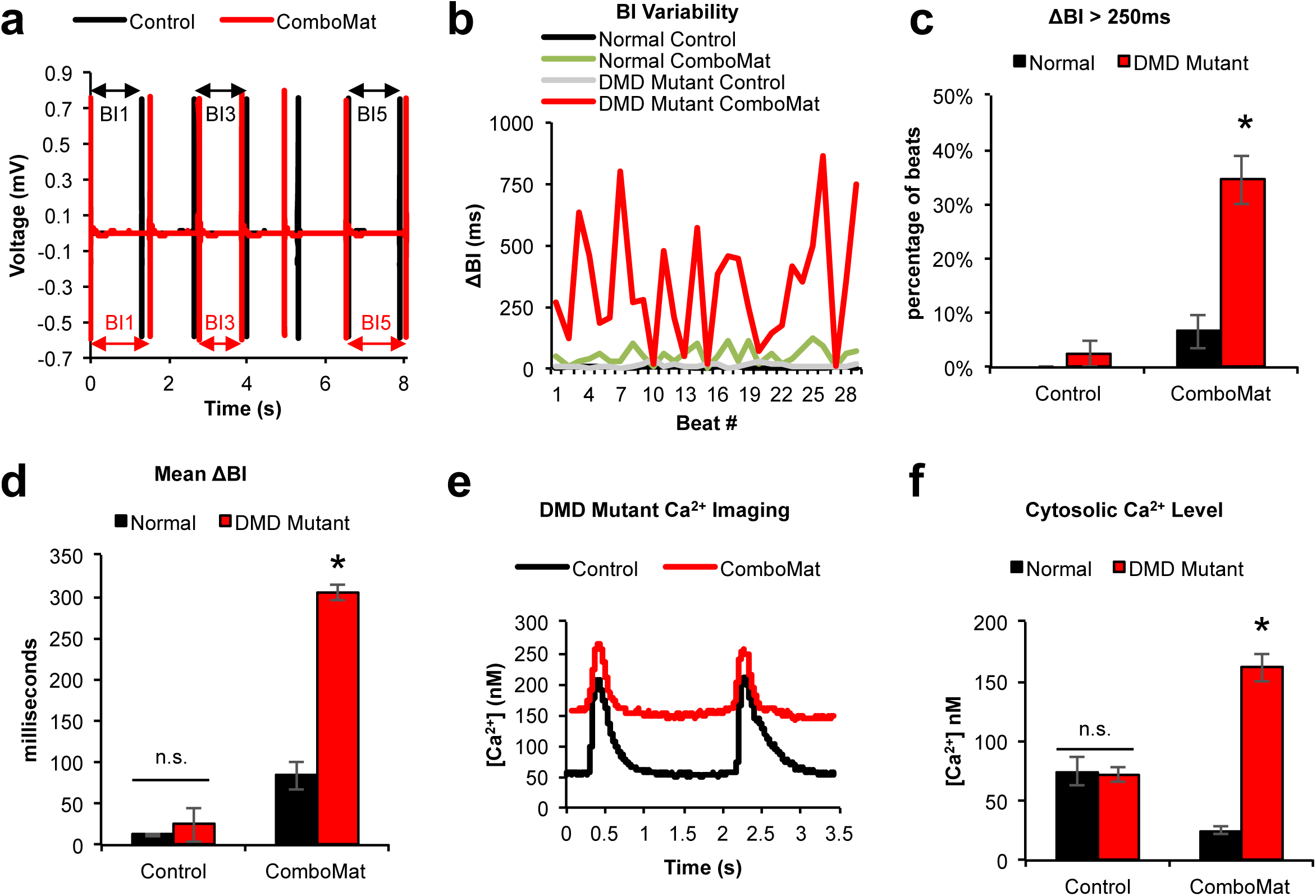
Endogenous DMD disease phenotype exposed. (**a**) Representative field potential recordings from *DMD* mutant hPSC-CMs exposed to Control (black) or ComboMat (red) conditions with every-other beat interval annotated. (**b**) Change in beat interval (ΔBI) plotted for 30 consecutive beats for Normal and *DMD* mutant hPSC-CMs exposed to Control or ComboMat conditions. Mature *DMD* mutant hPSC-CMs exhibit a much greater instability in ΔBI compared to Normal hPSC-CMs. (**c**) Average percentage of beats with a ΔBI > 250ms for Normal or *DMD* mutant hPSC-CMs exposed to Control or ComboMat conditions. Immature Normal hPSC-CMs showed no incidence of a ΔBI > 250ms. (**d**) The mean ΔBI for entire 30 beat recording. Immature *DMD* mutant hPSC-CMs show no distinguishing disease phenotype on the MEAs whereas *DMD* mutant hPSC-CMs matured using the ComboMat platform are significantly different from the Normal counterparts. (**e**) Representative Fura-2 Ca^2+^ imaging traces of *DMD* mutant hPSC-CMs exposed to Control (black) or ComboMat (red) conditions. (**f**) Quantification of diastolic Ca^2+^ content in the cytosol. *DMD* mutant hPSC-CMs exhibit a significantly higher cytosolic Ca^2+^ content when matured using the ComboMat platform. Bars represent averages ± S.E.M. *P<0.05.

To identify a potential mechanism for this beat rate variability, we measured intracellular calcium (Ca^2+)^ using the ratiometric Ca^2+^ indicator Fura-2 on an IonOptix imaging setup calibrated to calculate nanomolar intracellular Ca^2+^ concentrations. **Figure 5e** depicts representative Ca^2+^ traces from *DMD* mutant hPSC-CMs exposed to Control or ComboMat conditions. Matured *DMD* mutant hPSC-CMs were found to have an elevated baseline, or diastolic resting Ca^2+^ level, compared to the less mature cells in the Control group. Quantifying this diastolic Ca^2+^ level, we found a similar trend to the electrical instability presented previously. Normal and *DMD* mutant hPSC-CMs in the Control groups had cytosolic Ca^2+^ levels that were not statistically different from one another (**Fig. 5f**). *DMD* mutant hPSC-CMs matured using the ComboMat platform, however, have an elevated cytosolic Ca^2+^ concentration compared to Normal hPSC-CMs.

### Phenotypic drug screen using mature hPSC-CMs

In order to identify potential drug targets or classes of drugs for a more detailed phenotypic drug screen, we first performed a preliminary medium throughput, semi-automated screen with a small library of 2,000 diverse molecules with various mechanisms of action. Intracellular ATP (a widely used assay of cell viability in high throughput screens) was measured in *DMD* mutant hPSC-CMs after hypotonic stress. Overall the Z-prime of the assay was 0.71. The 8,000 data points (2,000 compounds at 4 concentrations) were ranked and plotted based on the Z-score (**Fig. 6a**). Taking into consideration the Z-score ranking (>3), standard deviation of replicates, and escalating dose-ranging response, 39 hits were identified from the 2000 input compounds (~2% hit rate) (**Supplementary Table 1**). Of these hits, 9 (~23%) were classified as Ca^2+^ channel blockers.

**Figure 6.**
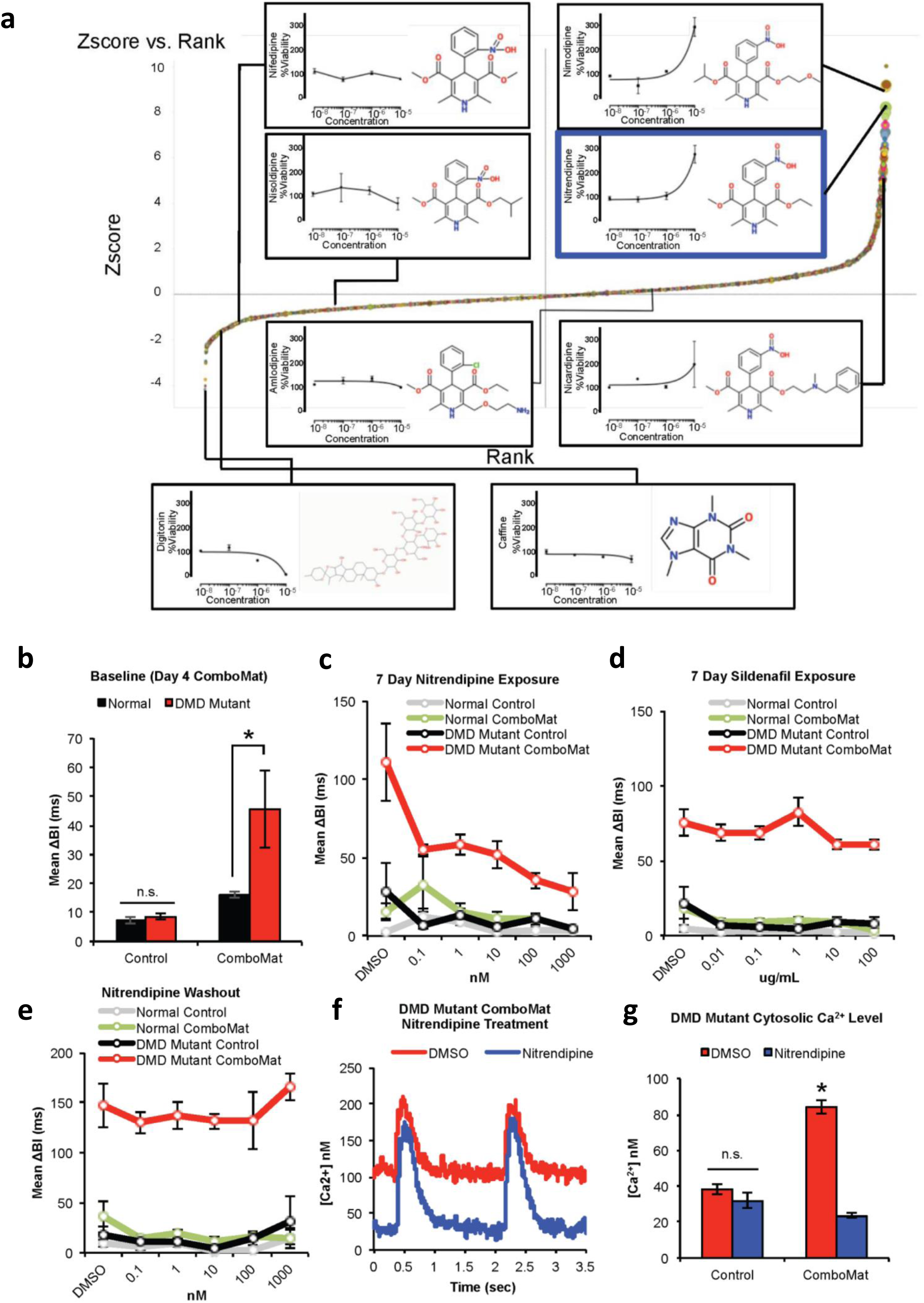
DMD cardiomyopathy phenotypic drug screen using mature hPSC-CMs. (**a**) Drug library ranked by Z-score for reduction in cell death in response to osmotic stress. Chemical structure of specific compounds also shown. (**b**) Quantification of baseline mean ΔBI for Normal and *DMD* mutant hPSC-CMs exposed to Control or ComboMat conditions. (**c,d**) Dose-response curve in response to nitrendipine and sildenafil after 7 days for Normal and *DMD* mutant hPSC-CMs exposed to Control or ComboMat conditions. (**e**) Measurement of mean ΔBI 48 hours after washout of nitrendipine. (**f**) Representative Fura-2 Ca^2+^ traces for mature *DMD* mutant hPSC-CMs and (**g**) quantification of diastolic Ca^2+^ content with (blue) and without (red) nitrendipine. Nitrendipine helps to reduce cytosolic Cal2+ load in the mature *DMD* mutant hPSC-CMs. Bars represent averages ± S.E.M. *P<0.05.

Incorporating this preliminary drug screen data with our findings of elevated intracellular Ca^2+^ content (**Fig. 5f**), we sought to utilize the nanoMEA platform to test whether nitrendipine (highlighted with blue box in **Fig. 6a**), a dihydropyridine Ca^2+^ channel blocker, could reduce the arrhythmogenic behavior of the mature *DMD* mutant hiPSC-CMs. We also tested sildenafil, an agent previously demonstrated not to benefit DMD patients^20^. Even at an early time point (four day ComboMat exposure) there was a significant difference in baseline beat rate variability between mature Normal hPSC-CMs and *DMD* mutant hPSC-CMs, while immature cells were indistinguishable from each other (**Fig. 6b**).

We then exposed cells to various concentrations of nitrendipine or sildenafil citrate for seven days and measured their beat rate variability. **Figure 6c,d** show the dose-response curves for nitrendipine and sildenafil, respectively. Nitrendipine significantly reduced the Mean ΔBI in mature *DMD* mutant hPSC-CMs compared to DMSO carrier control. Doses up to 1 µM nitrendipine restored beat rate variability to levels within physiologically relevant ranges (<50 ms). Higher doses, however, ceased spontaneous beating. In contrast, sildenafil had no effect on beat rate variability, regardless of the dose. Neither nitrendipine nor sildenafil had a significant impact on beat rate variability for cells exposed to Control conditions or Normal hPSC-CMs exposed to the ComboMat platform. The benefits of nitrendipine were washed out within 48 hours if removed from the cell culture medium (**Fig. 6e**). Ca^2+^ imaging revealed that 100 nM nitrendipine reduced the diastolic Ca^2+^ content in the mature *DMD* mutant hPSC-CMs compared to DMSO control (**Fig. 6f,g**). Nitrendipine had no impact on the diastolic Ca^2+^ content in cells exposed to the Control condition (**Fig. 6g**).

## DISCUSSION

The ComboMat platform proved to be a powerful tool in modulating cardiomyocyte development *in vitro*, increasing cell area, producing longer resting sarcomere lengths, and achieving more physiologically relevant binucleation percentages. Many of the functional enhancements that we observed from the ComboMat platform were predicted by our RNAseq analysis. GO terms for muscle development and contraction were upregulated in the ComboMat group, which resulted in faster contraction kinetics as measured by our CCQ contraction mapping. Metabolic GO terms such as fatty acid import were also upregulated in the ComboMat group and we found subsequently that cells exposed to the ComboMat group had a larger maximum respiratory capacity and ability to utilize exogenous fatty acids. Thus, there was agreement between the transcriptional and functional changes imparted by the ComboMat platform providing validation of our methods.

An important aspect of the present work is the development of a successful phenotypic drug screen platform. Methods such as the ComboMat platform, in conjunction with the nanoMEA electrophysiological screening system, can be scaled up for higher throughput screens of novel compounds. The noninvasive measurements can be used to monitor cell function longitudinally and the strong stratification of disease phenotype allows for easy comparison to healthy controls to see if drug treatment correlates with a restoration of function. We were able to validate that Ca^2+^ channel blockers were indeed a true hit from our initial screen as well as correctly weed out a false positive drug hit from *mdx* mouse model studies^20^.

Previous hPSC-CM models of dystrophic cardiomyopathy have relied heavily on hypotonic challenge to bring out a measurable disease phenotype^11^^,^^23^. While effective, hypotonic challenge is not physiologically accurate and is only a transient insult. Here we show that by maturing the *DMD* mutant hPSC-CMs with the ComboMat platform, cells present with beat rate variability due in part to calcium overload. Along with its involvement in mediating cell injury through reactive oxygen species (ROS) and mitochondrial apoptosis signaling pathways^27^, Ca^2+^ overload has long been known to mediate cardiac arrhythmias^28^ and has been implicated both *in vitro* and *in vivo* to be a driving force behind DMD pathophysiology^29^. Thus, we have demonstrated a maturation-dependent disease phenotype in *DMD* mutant hPSC-CMs that recreates an early disease phenotype found in patients. This is an important distinction from previous DMD disease models and one that we believe represents a logical progression in the cardiac disease modeling field. Wen et al. also reported the need for maturation in their hPSC-CM model of arrhythmogenic right ventricular dysplasia in order to bring out a disease phenotype^30^ and Tiburcy et al. demonstrated maturation via chronic catecholamine stimulation improved their ability to model heart failure^31^. Based on our findings, we propose that other cardiomyopathies may be more faithfully modeled by using hPSC-CMs matured beyond the typical fetal stage of standard directed differentiation methods.

While it is known that DMD patients exhibit diastolic dysfunction and arrhythmias, the pathogenesis of this disease phenotype is not fully understood^18^. Ca^2+^ overload has long been postulated to play a central role in the pathophysiology of dystrophic cardiomyopathy^29^. Leaky Ca^2+^ channels on the cell membrane and sarcoplasmic reticulum, abnormal NO signaling, and tears in the sarcolemma have all been proposed as the means for Ca^2+^ entry into the cell but no consensus exists. The present model, with an endogenous electrophysiological disease phenotype due to elevated Ca^2+^ levels, could prove to be a valuable tool for addressing some of these questions. Taken together, the present data show that cardiac maturation is an important factor in accurately modeling adult onset diseases, such as DMD cardiomyopathy, and that the ComboMat platform is essential in generating more mature hPSC-CMs. Furthermore, nanoMEA platform can be used in conjunction to effectively screen potential therapeutic targets.

## METHODS

### Experimental groups

RNAseq was performed on cells exposed to all different combinations of the three maturation cues: NPs, T3, and Let7i. We also included an empty vector (EV) negative control for virus exposure. For all other assays, we compared the full <u>ComboMat</u> platform (Let7i+NP+T3) to a viral vector <u>Control</u> (EV+Flat+No T3) and each maturation cue in isolation: <u>NP</u> (EV+NP+No T3), <u>T3</u> (EV+Flat+T3), and <u>Let7i</u> (Let7i+Flat+No T3). The ComboMat platform utilizes nanotopographic substrate cues, thyroid hormone T3, and Let-7i microRNA (miR) overexpression at specific time-points during the culture period to induce advanced structural and phenotypic development of hPSC-CMs. Each cue was chosen to impact specific aspects of hPSC-CM development. Nanotopographic substrate cues have been shown to promote the structural maturation of primary and hPSC-CMs^14^^,^^32^. These biophysical cues, however, have minimal impact on the metabolic development of hPSC-CMs. Therefore, thyroid hormone T3 was introduced as a biochemical cue to boost the metabolic, calcium handling, and contractile performance of the hPSC-CMs^16^. Preliminary results found, though, that long-term T3 exposure had a negative impact on sarcomere development and beat rate variability (**Supplementary Fig. 5**), reminiscent of complications from hyperthyroidism. To counteract these detrimental effects, we sought to manipulate the transcriptome using miRs such as the Let7 family. Along with improving metabolic capacity, hypertrophy, and the genetic profile^17^, Let7i helps to regulate the expression of the Hair Growth Associated gene that acts as a transcriptional co-repressor of many nuclear receptors, including the thyroid hormone receptor.

### hiPS cell line generation and maintenance

Normal urine-derived human induced pluripotent stem cells (hiPSCs) were obtained from an IRB-approved protocol as previously described^11^. Briefly, a polycistronic lentiviral vector encoding human Oct3/4, Sox2, Klf4, and c-Myc^33^ was used to reprogram cells collected from clean-catch urine samples into iPSCs^11^. The derivative hiPSC line (UC3-4) was karyotyped and shown to be normal 46, XY karyotype. hiPSCs were cultured on Matrigel-coated tissue culture plastic, fed mTeSR culture medium (StemCell Technologies), and passaged using 1.9 U/mL Dispase (ThermoFisher). Cells were cultured in hypoxic conditions at 37°C, 5% CO_2_, and 5% O_2_. *DMD* mutant hiPSCs (UC1015-6) were generated using CRISPR-Cas9 technology to create an isogenic pair from the Normal parental line UC3-4. Briefly, guide sequences targeting the first muscle specific exon of human dystrophin were used to delete a single G base at position 263 of the dystrophin gene to create a mutant allele (**Supplementary Fig. 1**). Mutation of the dystrophin gene was confirmed via sequencing. *DMD* mutant hiPSCs were maintained in the same manner as the Normal UC3-4 parental line.

### Cardiac directed differentiation and cell culture

A monolayer-based directed differentiation protocol was used as previously described^22^. hiPSCs were dissociated into single cells using Versene (ThermoFisher) and plated at a high density (250,000 cells/cm^2^) in mTeSR onto Matrigel-coated plates. Once a monolayer had formed, cells were treated with 1µM Chiron 99021 (Stemgent) in mTeSR until induction with 100nM Activin-A (R&D Systems) in RPMI 1640 (ThermoFisher) supplemented with B27 minus insulin (RPMI/B27/Ins-) and a switch to normoxic conditions (37°C, 5% CO_2_, ambient O_2_) at day 0. After 18 hours, cells were fed 5 ng/mL BMP4 (R&D Systems) plus 1µM Chiron 99021 in RPMI/B27/Ins- medium. At day 3 post-induction, cells were fed 1µM Xav 939 (Tocris) in RPMI/B27/Ins- medium. At day 5 post-induction, cells were fed fresh RPMI/B27/Ins- medium. At day 7 post-induction, cells were switched to RPMI-B27 medium containing insulin and medium (RPMI/B27/Ins+) was refreshed every other day. For T3 treated groups, 20 ng/ml T3 was freshly added to the RPMI/B27/Ins+.

### Lentiviral transduction of Let7i miRNA and purification of cardiomyocytes

The full description of the creation of the Let7i overexpression (OE) construct can be found in Kuppusamy et al.^17^. The control vector to Let7i was an eGFP expression pLKO.1 TRC vector (Addgene) under the U6 promoter. An eGFP construct was cloned between AgeI and EcroRI sites of the pLKO.1 TRC vector. To generate lentiviral particles, 293FT cells were plated one day before transfection. On the day of transfection, 293FT cells were co-transfected with psPAX2 (Addgene #12260), pMD2.G (Addgene #12259), vector and polyethylenimine (PEI). Medium was changed 24 hours later and the lentivirus particles were then harvested 24 and 48 hours later. Lentiviral particles were concentrated using PEG-it Virus Precipitation Solution (5x) (System Biosciences).

At day 15 post-induction, beating monolayers of hiPSC-CMs were dissociated with 10 U/mL collagenase type I in DPBS containing calcium and 10 U/mL DNAse for 1 hour at 37°C. Cells were then collected, spun down, and resuspended in TrypLE (ThermoFisher) containing 10 U/mL DNAse for 5 minutes. A P1000 micropipette was used to triturate clumps of hiPSC-CMs in the TrypLE until single cells were obtained. Cells were then plated onto Matrigel-coated plated at 100,000 cells/cm^2^. The following day, viral transduction of the cells was performed by diluting virus in the presence of hexadimethrine bormide (Polybrene, 6µg/ml) in RPMI/B27/Ins+ medium overnight. Cells were washed with PBS the following day and RPMI/B27/Ins+ was replaced.

Cardiomyocytes were then purified via metabolic challenge^34^ by feeding cells DMEM without glucose, sodium pyruvate, or glutamine (ThermoFisher) containing 2% horse serum and 4 mM lactate for three days without media change. Flow cytometry for cardiac purity was performed to confirm high purity samples (Supplementary Fig. 6). Afterward, hiPSC-CMs were switched back to RPMI/B27/Ins+ media containing 2 µg/mL puromycin dihydrochloride for three days to select for successfully transduced hiPSC-CMs.

### Fabrication of nanopatterned substrates and nanoMEAs

Nanopatterned substrates were fabricated via capillary force lithography as previously described^35^. Polyurethane acrylate (PUA) master molds were fabricated by drop dispensing liquid PUA pre-polymer onto a deep-ion etched silicon master with the intended nanotopographic geometries (800nm wide ridges and grooves, with 600nm depth) and placing a transparent polyester (PET) film on top. The nanopattern dimensions were chosen to enhance the structure and function of hPSC-CMs (**Supplementary Fig. 7**) After curing under a high-powered UV light source, the PUA and PET film were peeled from the silicon master to create a PUA master mold. To create the nanopatterned cell culture substrates, a polyurethane-based UV-curable polymer (NOA76, Norland Optical Adhesive) was drop dispensed onto glass primer treated 18mm, #1 coverslips (Fisher Scientific) and the PUA master mold was placed on top. After curing with a UV light source, the PUA master mold was peeled away, leaving behind the nanopatterned surface (NPs). Control flat substrates were fabricated using the same process but replacing the nanopatterned PUA master with an unpatterned PET film.

To prepare surfaces for cell culture, PU-based NPs and control flat substrates were washed with 70% ethanol, oxygen plasma treated at 50 W for 5 minutes, placed under a UV-C light source for 1 hour to sterilize, then incubated for 24 hours at 37°C with 5 µg/cm^2^ human fibronectin (Life Technologies). hPSC-CMs were plated at a density of 150,000 cells/cm^2^. Nanopatterned MEAs (nanoMEAs) were fabricated as described [Smith, A.S.T. *et al*. Nanotopographically-patterned microelectrode arrays for high-throughput analysis of human cardiac structure-function relationships. *PNAS*. Under Review]. Briefly, the ion permeable polymer Nafion (Sigma-Aldrich) was patterned onto commercial multi-well MEA plates (Axion Biosystems) via solvent-mediated capillary force lithography. Prior to cell seeding, control flat or nanoMEA Nafion surfaces were sterilized with UV-C light and then incubated for 6 hours at 37°C with 5 µg/cm^2^ human fibronectin (Life Technologies). hPSC-CMs were plated in droplet fashion at 25,000 cells per well of the MEA plate.

### Immunofluorescent imaging and morphological analysis

After three weeks of the ComboMat platform, cells were prepared for immunofluorescent analysis by fixing in 4% paraformaldehyde (Affymetrix) for 15 minutes at room temperature. Samples were then permeabilized with 0.1% Triton X-100 (Sigma-Aldrich) in PBS and blocked with 5% goat serum. Cells were then incubated with mouse anti-α-actinin monoclonal antibody (1:1000, Sigma-Aldrich) in 1% goat serum overnight at 4°C. After washing with PBS, the samples were stained with Alexa-594 conjugated goat-anti-mouse secondary antibody (1:200, Invitrogen) as well as Alexafluo 488 labeled Phalloidin (1:200, Invitrogen). Nuclei counterstaining was performed with Vectashield containing DAPI (Vector Labs). A Nikon A1R confocal microscope with a Nikon CFI Plan Apo VC 60x water immersion objective was used to capture detailed immunofluorescent images for sarcomere analysis while a 20x air objective was used to obtain low-powered images of cells for measuring cell binucleation percentage. Cells from both objective powers were used to measure cell size. For morphological and sarcomere analysis, hPSC-CMs were plated at 10,000 cells/cm^2^. For each condition, at least three biological specimens were plated with at least 7 cells imaged per specimen.

### Quantitative sarcomere analysis

Immunofluorescent images were quantified for sarcomere alignment, pattern strength, and sarcomere length using a scanning gradient / Fourier transform script in MATLAB (Mathworks). Each image was broken into several small segments which were individually analyzed. Using a directional derivative, the image gradient for each segment was calculated to determine the local alignment of sarcomeres. The pattern strength was then determined by calculating the maximum peaks of one-dimensional Fourier transforms in the direction of the gradient. The lengths of sarcomeres were calculated by measuring the intensity profiles of the sarcomeres along this same gradient direction. The frequency at which the intensity profiles crossed their mean allowed an accurate calculation of local sarcomere length within each image segment. Once completed, this analysis allowed for unbiased subcellular-resolution mapping of sarcomere patterning, even in cells lacking the alignment cues provided by nanotopography.

### Calcium imaging

Intracellular calcium content was measured using the ratiometric indicator dye fura2-AM as described previously^36^. In brief, cells were incubated in 0.2µM fura2-AM dye in Tyrode’s solution for 20 minutes at 37°C and washed with PBS. Spontaneous Ca^2+^ transients were then monitored with the Ionoptix Stepper Switch system coupled to a Nikon inverted fluorescence microscope. The fluorescence signal was acquired using a 40x Olympus objective and passed through a 510 nm filter, and the signal was quantified using a photomultiplier tube. Diastolic resting calcium concentration was quantified in the IonOptix software IonWizard using the formula:

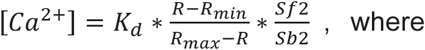

Kd = fura2 calcium dissociation constant = 225nM

R_max_ and R_min_ = ratio values measured under saturating calcium levels and in the absence of calcium, respectively.

Sf2/Sb2 = ratio between the background subtracted wavelength 2 excited fluorescence in zero and saturating calcium solutions, respectively.

### Electron microscopy and ultrastructural analysis

After three weeks under the ComboMat platform, hiPSC-CMs were fixed in 4% glutaraldehyde in sodium cacodylate buffer for 2 hours at room temperature for transmission electron microscopy (TEM) analysis. Samples were then washed in buffer and stained in buffered 1% osmium tetroxide for 30 minutes on ice. Next, samples were washed in water and dehydrated in a graded series of ethanol. Samples were then treated by infiltration of 1:1 ethanol:Epon-Araldite epoxy resin followed by two changes of pure Epon-Araldite. At this point, a second coverslip was placed on top of the samples in a 60°C oven for polymerization. The following day, the coverslips were dissolved using hydrofluoric acid and the samples were mounted onto blocks for sectioning. Ultra-thin (70-80 nm) axial sections of single cells were contrasted with lead citrate prior to imaging with a JEOL JEM 1200EXII at 80 kV. ImageJ (National Institutes of Health) was used to measure Z-band width as defined as the width of contiguous electron-dense bands within a sarcomere structure.

### MEA electrophysiological analysis

Electrophysiological analysis of spontaneously beating cardiomyocytes was collected for 2 minutes using the AxIS software (Axion Biosystems). After raw data collection, the signal was filtered using a Butterworth band-pass filter and a 90 µV spike detection threshold. Field potential duration was automatically determined using a polynomial fit T-wave detected algorithm. Beat interval analysis on 30 consecutive beats was performed as previously published^26^ (see **Supplemental Table 2**).

### Mitochondrial functional assay

Cellular metabolic function was measured using the Seahorse XF96 extracellular flux analyzer as previously published^17^. After three weeks of treatment with the ComboMat platform, cells were trypsinized and replated onto fibronectin coated (5 µg/cm^2^) XF96 plates at a density of 30,000 cells per well. The MitoStress and Palmitate assays were performed three days post-replating onto the XF96 plates. One hour prior to experiments, RPMI/B27/Ins+ media was replaced with Seahorse XF Assay media supplemented with sodium pyruvate (Gibco/Invitrogen, 1mM) and 25mM glucose for the MitoStress assay or 25mM glucose with 0.5mM Carnitine for the Palmitate assay. For the MitoStress assay, injection of substrates and inhibitors were applied during the measurements to achieve final concentrations of 4-(trifluoromethoxy) phenylhydrazone at 1 μM (FCCP; Seahorse Biosciences), oligomycin (2.5μM), antimycin (2.5μM) and rotenone (2.5μM). For the Palmitate assay, injection of substrates and inhibitors were applied during the measurements to achieve final concentrations of 200mM palmitate or 33μM BSA. The OCR values were further normalized to the number of cells present in each well, quantified by the Hoechst staining (Hoechst 33342; Sigma-Aldrich) as measured using fluorescence at 355 nm excitation and 460 nm emission. Basal respiration in the MitoStress assay was defined as the OCR prior to oligomycin addition while maximum respiratory capacity was defined as the change in OCR in response to FCCP from the OCR after the addition of oligomycin. Exogenous fatty acid utilization was defined as the maximum change in OCR from baseline after the addition of Palmitate.

### Contraction analysis

Our video-based contraction analysis algorithm, termed Correlation-based Contraction Quantification (CCQ), utilizes particle image velocimetry and digital image correlation algorithms^37^ to track the motion of cells from bright field video recordings as previously published^22^. Functional outputs such as contraction magnitude, velocity, and angle are output as a vector field for each video frame and can be averaged spatially or temporally. In brief, a reference frame is divided into a grid of windows of a set size. To track motion, each window is run through a correlation scheme comparing a second frame, providing the location of that window in the second frame. The correlation equation used provided a Gaussian correlation peak with a probabilistic nature that provides sub-pixel accuracy. The videos used for this analysis were taken with a Nikon camera at 20x magnification and 30 frames per second.

### RNA-seq analysis

Total RNA was extracted using trizol (Thermo Fisher Scientific) from RUES2 hESC-CMs. RNAseq samples were aligned to hg19 using Tophat (PMID: 19289445, version 2.0.13). Gene-level read counts were quantified using htseq-count (PMID: 25260700) using Ensembl GRCh37 gene annotations. Genes with total expression above 3 RPKM summed across RNA-seq samples were kept for further analysis. *princomp* function from R was used to for Principal Component Analysis. DESeq (PMID: 20979621) was used for differential gene expression analysis. Genes with fold change >1.5 were considered differentially expressed. topGO R package (PMID: 16606683) was used for Gene Ontology enrichment analysis.

For the heat map of pathway enrichment, each condition was compared against the Control (EV-Flat) condition, and up-regulated genes (>1.5 fold change) and down-regulated genes were identified (< -1.5 fold change). Hypergeometric test was performed on up- and down-regulated genes separately for enrichment against a curated set of pathways that are beneficial for cardiac maturation, resulting in a m by n matrix, where m is the number of pathways (m=7) and n is the number of conditions (n=10). The negative log10 of the ratio between enrichment p-value for up- and down-regulated genes were calculated to represent the overall net “benefit” of a treatment: large positive value (>0) means the treatment results in more up-regulation of genes in cardiac maturation pathways than down-regulation of these genes, and more negative values means the treatment results in more down-regulation of genes in cardiac maturation pathways.

### Flow cytometry

Cells were dissociated and prepared for flow cytometry before and after lactate purification and puromycin selection to determine selection efficiency. Cells were first fixed in 4% paraformaldehyde for 15 minutes. Cells were then permeabilized with 0.75% Saponin and stained in PBS containing 5% FBS with either 1:100 mouse anti-cTnT or isotype control mouse IgG antibodies. Alexa-488 conjugated goat-anti-mouse secondary antibody (1:200, Invitrogen) was used for visualization. Samples were run on a BD Sciences FACS Canto flow cytometer and analyzed with FlowJo software.

### High-throughput drug screening

Cardiomyocytes were dissociated 14 days post differentiation into a single cell suspension by 5 minute incubation in TrypLE (Life Technologies) and plated at 10,000 cells per well on opaque-bottom 384-well plates (Nunc) pre-coated with 1 mg/mL Matrigel (Corning) for one hour at 37°C. Cells were then cultured on the 384-well plates for an additional 16 days, with media exchanges every 72 hours. After 15 days, compounds dissolved in DMSO were distributed to the plates using the CyBi Well Vario 384/25 liquid handler (Cybio, Germany) to achieve concentrations of 10^-8^, 10^-7^, 10^-6^, 10^-5^ M in duplicate. After 18 hours of incubation, medium was aspirated and pure water was injected to reach 12.5% normal osmolarity using BioTek EL406 Washer Dispenser (BioTek, Winooski, VT). After 30 minutes incubation, supernatants were removed and plates were assayed by adding CellTiter Glo (Promega) according to the manufacturer’s instructions. After 5 minutes, the luminescence was measured using EnVision Multilable Resaer (PerkinElmer). All data were processed and visualized by Tibco Spotfire (Tibco Spotfire, Boston, MA). Percentage viability was calculated by comparing signal from each well to the average of the control wells treated with DMSO alone (32 control wells per plate).

### Statistical analyses

Statistical significance between groups was determined using either a One-Way or Two-Way ANOVA with Tukey’s pairwise *post hoc* analysis from SigmaPlot software unless otherwise stated. For all statistical analyses, a p-value less than 0.05 was considered significant. Error bars represent standard error of the mean (SEM).

## ACKNOWLEDGEMENTS

This work is supported by grants from the National Institutes of Health R01 HL135143 (DHK), R01GM097372, and R01GM083867 (HRB), 1P01GM081619 (CM and HRB), and the NHLBI Progenitor Cell Biology Consortium U01HL099997 and UO1HL099993 (CM and HRB). The Vision Core for their TEM services P30 EY001730.

## AUTHOR CONTRIBUTIONS

DHK, HRB and JRM conceived and designed the study. JRM and JWM generated and analyzed all the experimental data with assistance from WL, XG and SL. DM, ASTS, YW, MR and MKC contributed to the analysis and interpretation of results and preparation of the manuscript. JRM, JWM, HRB and DHK and all other authors contributed to the writing of the paper.

